# The Effect of Adrenalectomy on Bleomycin-Induced Pulmonary Fibrosis in Mice

**DOI:** 10.1101/2024.01.31.577771

**Authors:** John McGovern, Carrighan Perry, Alexander Ghincea, Shuai Shao, Erica L. Herzog, Huanxing Sun

## Abstract

Progressive lung fibrosis is often fatal and has limited treatment options. Though the mechanisms are poorly understood, fibrosis is increasingly linked with catecholamines such as adrenaline (AD) and noradrenaline (NA), and hormones such as aldosterone (ALD). The essential functions of adrenal glands include the production of catecholamines and numerous hormones, but the contribution of adrenal glands to lung fibrosis remains less well studied. Here, we characterized the impact of surgical adrenal ablation in the bleomycin model of lung fibrosis. Wild type mice underwent surgical adrenalectomy or sham surgery followed by bleomycin administration. We found that the bleomycin induced collagen over deposition in the lung was not affected by adrenalectomy. However, histologic indices of lung remodeling were ameliorated by adrenalectomy. These findings were accompanied by a decrease in bronchoalveolar lavage (BAL) cell count along with concomitant reductions in alpha smooth muscle actin (⍺SMA) and fibronectin. Surgical adrenalectomy completely abrogated AD detection in all compartments, but only reduced NA in the BAL of uninjured mice. Systemic ALD levels were reduced after adrenalectomy. Taken together, these results support the presence of pulmonary-adrenal axis in lung fibrosis and suggest that adrenalectomy is protective in this disease. Further investigation will be needed to better understand this observation and aid in the development of novel therapeutic strategies.

## INTRODUCTION

Progressive pulmonary fibrosis (PPF) is characterized by progressive scarring of the lungs [1], remodeling of the lung’s extracellular matrix (ECM), and increased deposition of collagen and alpha-smooth muscle actin (αSMA) [2]. This is thought to be driven by elevated levels of fibroblasts that have differentiated into activated myofibroblasts [3, 4]. PPF is a devastating form of interstitial lung disease (ILD), has limited treatment options, and is often fatal, especially in idiopathic pulmonary fibrosis (IPF) with a life expectancy of 3-5 years after diagnosis [5]. Furthermore, the mechanisms behind these diseases are poorly understood, making it difficult to develop effective treatments [1]. A better understanding of the drivers of fibroblast to myofibroblast differentiation, collagen deposition, and ECM remodeling, and how these processes might be reversed, is paramount to the development of more effective therapies.

The adrenal glands are important regulators of systemic homeostasis and respond to various stimuli via their production of catecholamines and aldosterone (ALD). Emerging studies suggest that fibrosis is associated with catecholamines such as adrenaline (AD) and noradrenaline (NA) [6–10]. Additionally, work from our lab has shown that NA derived from lung-specific adrenergic nerves contributes to pulmonary fibrosis [6, 8, 11] which was ameliorated with alpha-adrenergic receptor 1D (ADRA1D) antagonism [8]. ALD, as the final effector of the renin angiotensin aldosterone system (RAAS), contributes to the regulation of ECM deposition and the activity of fibrotic mediators such as TGFβ1 [12]. A better understanding of these associations has the potential to provide insights into the mechanism(s) by which the adrenal glands and catecholamines contribute to pulmonary fibrosis and unveil new opportunities for treatment.

This study sought to address these knowledge gaps by assessing whether adrenalectomy might mitigate lung inflammation, biochemical collagen accumulation, lung histology, ECM remodeling, and catecholamine and ALD levels in the bleomycin model of experimental pulmonary fibrosis.

## METHODS

### Animals

All experiments involving animals were approved by Yale School of Medicine IACUC in accordance with federal regulations (protocol #20292). All experiments used C57 Black 6 (C57BL/6) wild-type mice. The mice underwent either an adrenalectomy or sham surgery (Figure 1) conducted by both our own lab personnel or by the Jackson Laboratory. The mice were injected subcutaneously with buprenorphine proximate to the time of the surgery. More than 10 minutes after the injection, the mice were administered one dose of ketamine and xylazine. A one-third re-dose was administered if proper sedation, assessed by absence of response to toe-pinch, was not yet achieved. The anesthetized mice were placed in ventral recumbency. Isoflurane was administered nasally by vaporizer and eye lubricant applied to both eyes. The mice were shaved from the hip to the mid-thorax on the dorsal side then lidocaine injected intradermally around the planned incisions. The skin was sterilely prepared with betadine and 70% ethanol. A 1-2 cm incision, adjusted to the size of the animal, was made in the mid-dorsal area, with its extreme cranial end at the level of the 13^th^ rib. A window through the muscle fibers was made by entering the fibers and separating them bluntly with sharp (iris) scissors. The adrenal gland was removed intact with forceps without any cutting. The muscle layer was closed with absorbable suture and the skin was closed with surgical clips. Immediately after surgery, the mice were subcutaneously injected with meloxicam and 500 µL of 0.9% NaCl. Meloxicam was injected every 24 hours for three days and 1% saline was added to the drinking water to counteract mineralocorticoid deficiency. After 7 days, the surgical clips were removed under anesthesia. After the mice were allowed to recover from the surgery for three weeks, they were administered either a single dose of 1.5 U/kg pharmacologic grade bleomycin (Northstar Rx LLC, NDC 16714-8860-0) or sterile saline, by orotracheal aspiration [13]. Mice were anesthetized with isoflurane and suspended by their incisors on a standing rack. With the tongue held in gentle retraction, 50 µL bleomycin or saline was pipetted into the oropharynx and aspirated. All mice were humanely sacrificed 21 days after for sample collection.

**Figure 1.**
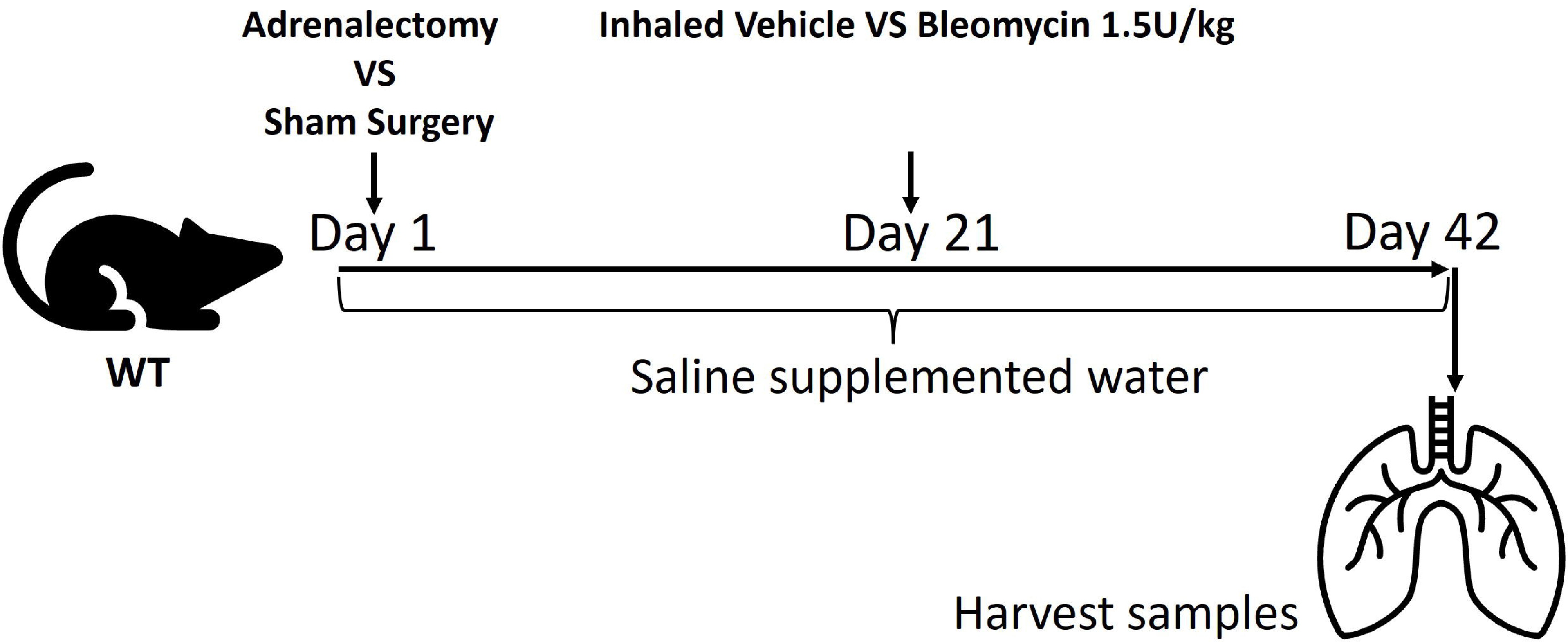
Adrenalectomy and sham surgery were applied to bleomycin model. WT mice underwent surgical adrenalectomy, or sham surgery. After 21 days mice were anesthetized with isoflurane, and bleomycin (1.5 U/kg of body weight) dissolved in saline or vehicle control were administered orotracheally as a single dose. The samples were harvested at 42 days.

### Bronchoalveolar Lavage (BAL) Cell Quantification

At the time of sacrifice, two aliquots of 0.8 mL of PBS were slowly instilled into lung and the lavage fluid gently aspirated. The combined BAL sample was assessed for white blood cell counts using a Beckman Coulter Ac.T Diff instrument.

### Soluble and Insoluble Collagen Quantification

The soluble and insoluble collagen concentrations of the three lobes (superior, middle, and inferior lobes) of the right lungs of the sacrificed mice were quantified using the Sircol Soluble Collagen Assay Kit (Biocolor Ltd., CLS 1111) and the Sircol Insoluble Collagen Assay Kit (Biocolor Ltd., CLS 2000), following manufacturer instructions.

### Trichrome Staining and Scoring

Whole left lungs and the post-caval lobes were formalin fixed and paraffin embedded (FFPE) and sectioned for histologic analysis with Masson’s trichrome stain to assess collagen deposition. Six images per slide were taken at 20X magnification and assigned a modified Ashcroft score (MAS), as previously described [14]. The scores were then averaged for each slide to determine an overall score for each lung.

### Immunofluorescent (IF) Staining and Analysis

The FFPE sectioned left lungs were also used to assess for ECM changes resulting from adrenalectomy. The slides were de-paraffinized in xylene and rehydrated in decreasing concentrations of ethanol in water. Antigen retrieval was done using BD Pharmingen Retrievagen A (Cat# 550524), following manufacturer instructions. The sections were penetrated with PBS+0.25% Triton and blocked for 30 minutes using 3% bovine serum albumin (BSA) in PBS. Tissues were stained to test alpha smooth muscle actin (αSMA) (Abcam, Cat# ab7817), laminin (Abcam, Cat# ab11575) and fibronectin (Rabbit pAb Abcam, Cat# ab23751). All antibodies were diluted 1:250 in 1% BSA in PBS. Slides were incubated overnight at 4°C and stained with secondary antibody (Alexa Fluor 555 donkey anti-rabbit IgG (H+L) Invitrogen, Cat# A31572 and Alexa Fluor 488 chicken anti-mouse IgG (H+L) Invitrogen, Cat# A21200), diluted 1:500 in 1% BSA in PBS. The samples were incubated for one hour at 37°C then mounted using VECTASHIELD Mounting Media with DAPI and covered (Cat# H-1200).

Images were taken at 20X and 40X magnification using a Nikon Eclipse Ti microscope and the NIS Elements Br software. Images were merged and analyzed using ImageJ v1.54f. A threshold was set manually to separate positive from negative signals. The threshold was applied to all images taken from the same channel. Mean fluorescent intensity (MFI) and the expression areas of αSMA, laminin and fibronectin were individually calculated. The JACoP plugin was used to measure the overlapping area of laminin or fibronectin with αSMA [15].

### Noradrenaline and Adrenaline Quantification by ELISA

NA and AD concentrations from plasma, BAL, and right lung homogenates were quantified by ELISA using the Noradrenaline Research ELISA kit (LDN, BA E-5200R) and the Adrenaline Research ELISA kit (LDN, BA E-5100R), following manufacturer instructions.

### Aldosterone Quantification by ELISA

Plasma aldosterone concentration was measured using an Aldosterone ELISA kit (LDN, MS E-5200R), following manufacturer instructions. All plasma samples were diluted 1:10 with the Standard A provided with the kit (LDN, E-5201).

### Statistical Analyses

Statistical analysis was performed using GraphPad Prism version 10.0.2. For parametric comparisons of two conditions, we used an unpaired Student’s t test with Welch’s correction. For non-parametric comparisons of more than two conditions, we used ordinary one-way ANOVA with multiple comparisons, comparing the mean of a column to the mean of every other column. We considered a P value less than 0.05 to be significantly different.

## RESULTS

### Adrenalectomy is protective against bleomycin-induced lung inflammation

Adrenally derived hormones including NA, AD, and ALD are associated with inflammation [16–19]. Reasoning that adrenalectomy would lead to a decrease in these hormone levels and therefore improved markers of inflammation, we evaluated BAL white blood cell (WBC) counts. Consistent with former studies [20], we found that bleomycin results in significantly increased airway inflammation by BAL WBC count, and that this increase was ameliorated by adrenalectomy (Figure 2). This indicates that adrenalectomy is protective against airway inflammation in the bleomycin model of pulmonary fibrosis.

**Figure 2.**
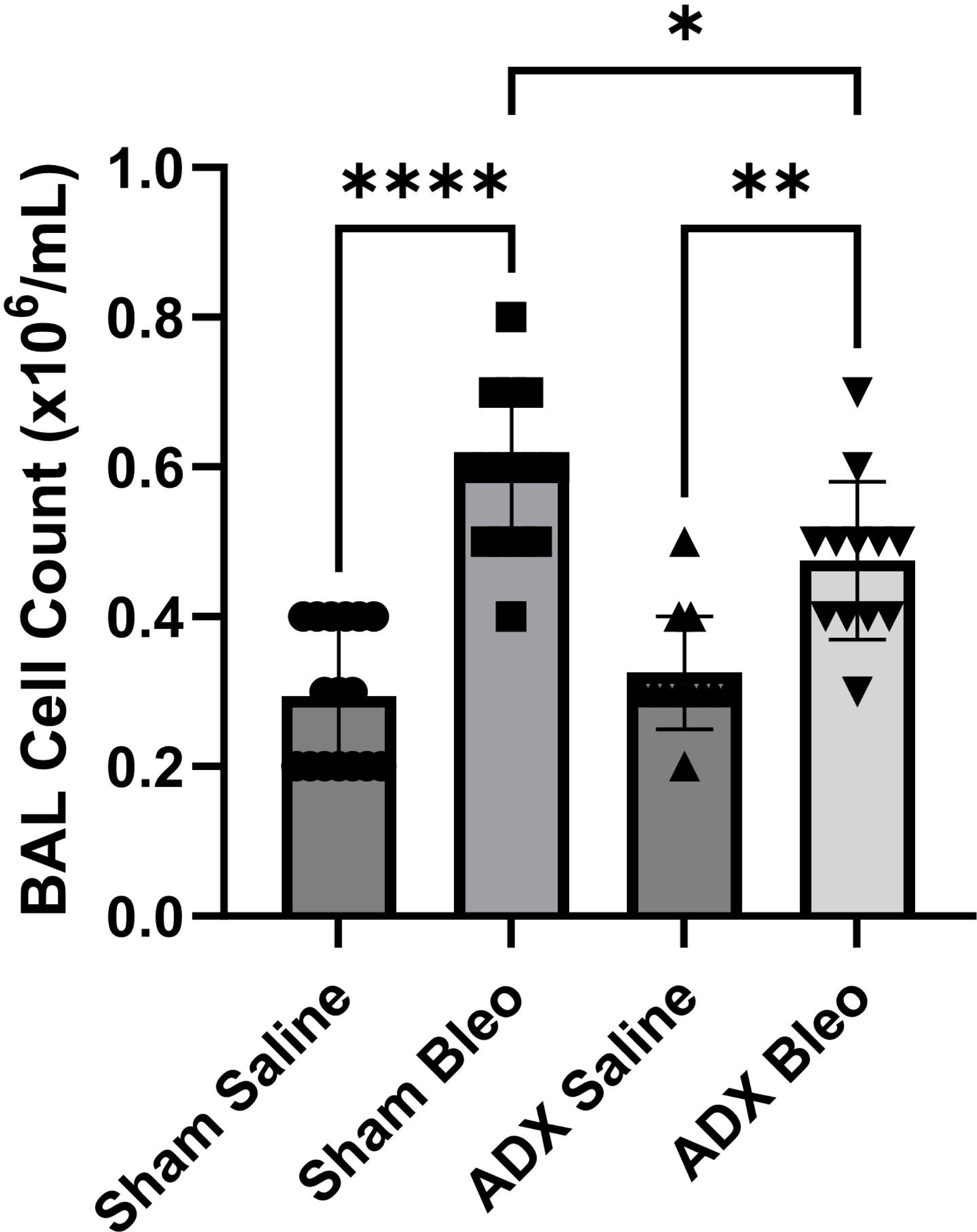
Adrenalectomy Inhibits lung inflammation in the bleomycin model. Relative to sham surgery, adrenalectomy has significant impact on lung inflammation in the bleomycin model as measured by BAL cell counts. Data are shown as mean +/- SEM. One-way ANOVA was performed to determine significance (* = p<0.05; ** = p<0.01; *** = p<0.005; **** = p<0.0001).

### Adrenalectomy mitigates bleomycin-induced fibrotic remodeling

Work by our lab and others has shown that catecholamines and aldosterone contribute to the development of tissue fibrosis [6, 8, 21]. We therefore reasoned that adrenalectomy may have an effect on tissue remodeling and fibrosis. To investigate this question, we examined Masson’s trichrome stained FFPE lung tissue sections obtained from bleomycin-treated mice. We found that mice subjected to adrenalectomy had a significant decrease in MAS when compared to the sham surgery group (Figure 3). This suggests that adrenalectomy may have a protective effect on mice with bleomycin induced lung fibrosis.

**Figure 3:**
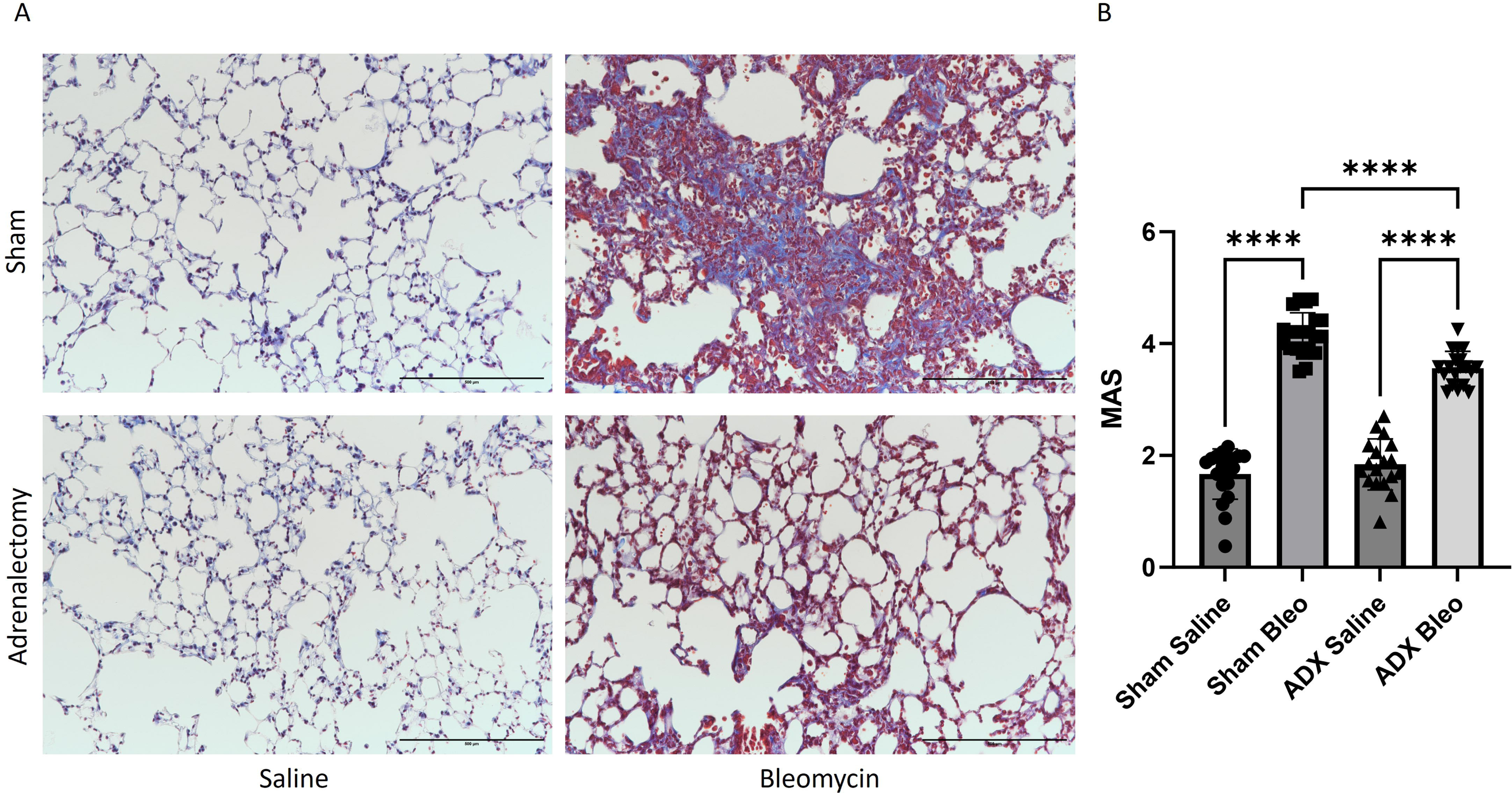
Adrenalectomy impacts Bleomycin-induced lung remodeling. (A) In the bleomycin model, trichrome staining to evaluate the lungs of mice that underwent adrenalectomy vs sham surgery. (B) After treatment with Bleomycin, a significant increase in the Modified Ashcroft Score (MAS) was observed in lung tissues of mice from both the adrenalectomy and sham surgery groups. Notably, adrenalectomy resulted in a significant reduction of MAS when compared to the sham-operated counterparts in mice that received Bleomycin. Scale bar = 500 microns. Data are shown as mean +/- SEM. One-way ANOVA was performed to determine significance (**** = p<0.0001).

### Adrenalectomy does not reduce collagen deposition

In order to further examine mechanisms underlying adrenalectomy’s protective effects in the bleomycin model, we next decided to quantify the soluble and insoluble collagen content of lung tissues as these measurements provide a biochemical surrogate for the severity of lung fibrosis. Using the Sircol assay we found that there was no change in either the soluble or insoluble collagen between the sham and adrenalectomy groups in bleomycin-treated mice (Figure 4). Interestingly though, there was an increase in both soluble and insoluble collagen in control mice after adrenalectomy. Together, these results indicate that adrenalectomy does not reduce collagen deposition in the bleomycin model of pulmonary fibrosis and may actually lead to increases in overall collagen content. Thus, the improvements in fibrosis observed by histopathologic examination are likely to be the result of changes in the non-collagen components of the ECM.

**Figure 4:**
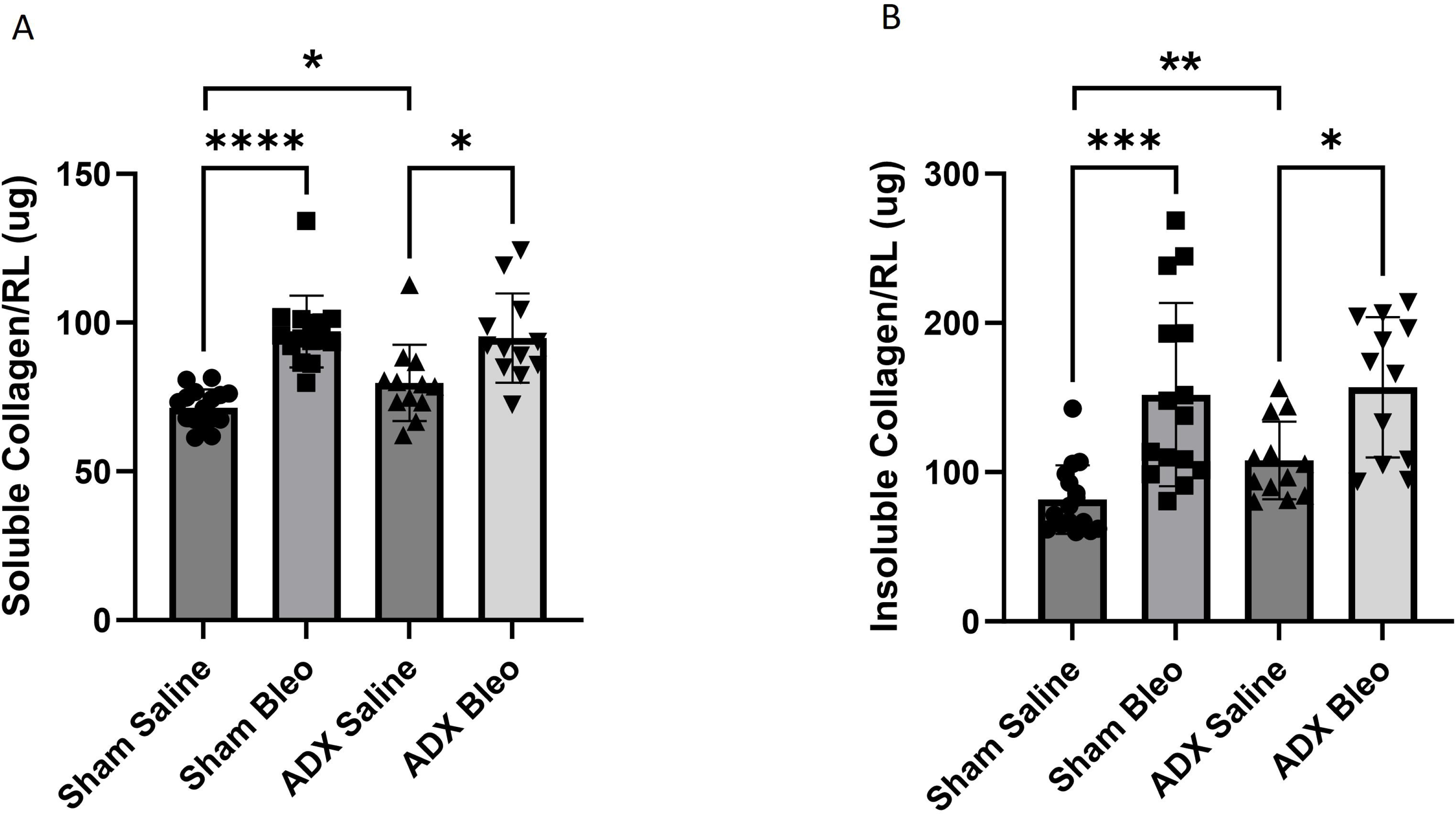
The effect of Adrenalectomy on lung collagen accumulation. After treatment with Bleomycin, both soluble (A) and insoluble (B) collagen were measured in the right lung tissues of mice from both the adrenalectomy and sham surgery groups. Notably, adrenalectomy resulted in significant reductions of both soluble (A) and insoluble (B) collagens when compared to the sham surgery counterparts in the mice that received vehicle control (saline), but not Bleomycin. Data are shown as mean +/- SEM. One-way ANOVA was performed to determine significance (**** = p<0.0001).

### Adrenalectomy reduces αSMA expression

To better understand the mechanisms behind our observation that surgical ablation of the adrenal glands improves histopathologic markers of fibrosis, we next investigated whether other fibrotic markers were also affected by adrenalectomy. ⍺SMA expressing myofibroblasts are well-known to accumulate in fibrotic regions of the lung and contribute to the development and progression of pulmonary fibrosis [22]. Hence, we first evaluated the expression of αSMA in lung tissues by IF staining. Using this method, we observed a significant reduction in both the expression area and mean fluorescent intensity (MFI) of αSMA in the adrenalectomy/bleomycin groups compared to the sham/bleomycin group indicating that adrenalectomy induces a reduction in ⍺SMA expression (Figure 5).

**Figure 5:**
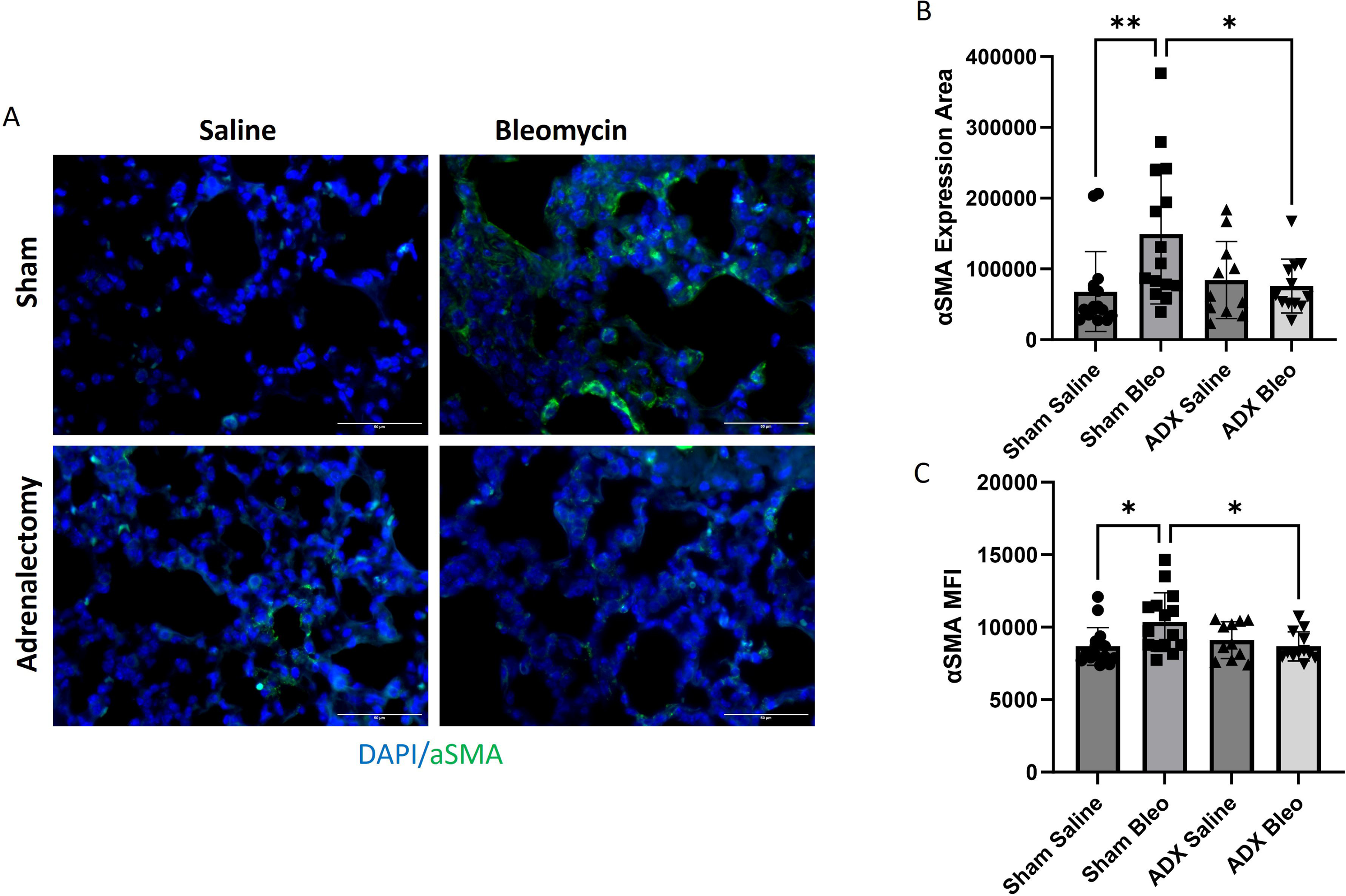
The effect of Adrenalectomy on αSMA expression in Bleomycin challenged mice lung. (A) Immunostaining of α-smooth muscle actin (αSMA) in the lung tissue of mice that underwent adrenalectomy or sham surgery in the Bleomycin model of lung fibrosis. For all images αSMA is in the green, FITC channel, and nuclei are counterstained with DAPI. Bleomycin treatment significantly increased both (B) the αSMA positive area and (C) the expression intensity (Mean Fluorescence Intensity, MFI) of αSMA. This was significantly reversed by adrenalectomy. Scale bar = 50 microns. Data are shown as mean +/- SEM. One-way ANOVA was performed to determine significance (* = p<0.05; ** = p<0.01).

### Adrenalectomy is protective against fibrotic ECM remodeling

Besides collagen, there are other core ECM proteins associated with fibrotic disease, such as laminin and fibronectin [23–26]. In order to test the hypothesis that adrenalectomy improves remodeling in the lung we also evaluated these ECM components using IF staining. We first measured the expression of laminin and observed a small but statistically significant increase in laminin expression by MFI in sham mice that received bleomycin (Figure 6A and B). Adrenalectomy eliminated this bleomycin-induced augmentation in laminin MFI though did not otherwise decrease laminin expression (Figure 6B and C). Thus, while adrenalectomy does not decrease overall laminin expression, it does appear to offer some protection against laminin associated fibrotic remodeling in the bleomycin model of lung fibrosis. To further verify this observation, we measured the co-expression of laminin and αSMA and found that there was significantly less overlap in the expression of laminin and ⍺SMA in bleomycin-treated mice subjected to adrenalectomy as opposed to sham surgery (Figure 6A and D). Turning to fibronectin, we discovered that adrenalectomy significantly mitigates increases in fibronectin expression caused by bleomycin as determined by MFI and area of expression (Figure 7A-C). As with laminin, we also assessed the co-expression of fibronectin and ⍺SMA and found that adrenalectomy effectively protects against increases in fibronectin and ⍺SMA co-expression that are induced by bleomycin (Figure 7D). Taken together, these results indicate that adrenalectomy protects against fibrotic remodeling of the pulmonary ECM in the bleomycin model by mitigating increases in laminin and fibronectin expression and by reducing the co-expression of these glycoproteins with ⍺SMA.

**Figure 6:**
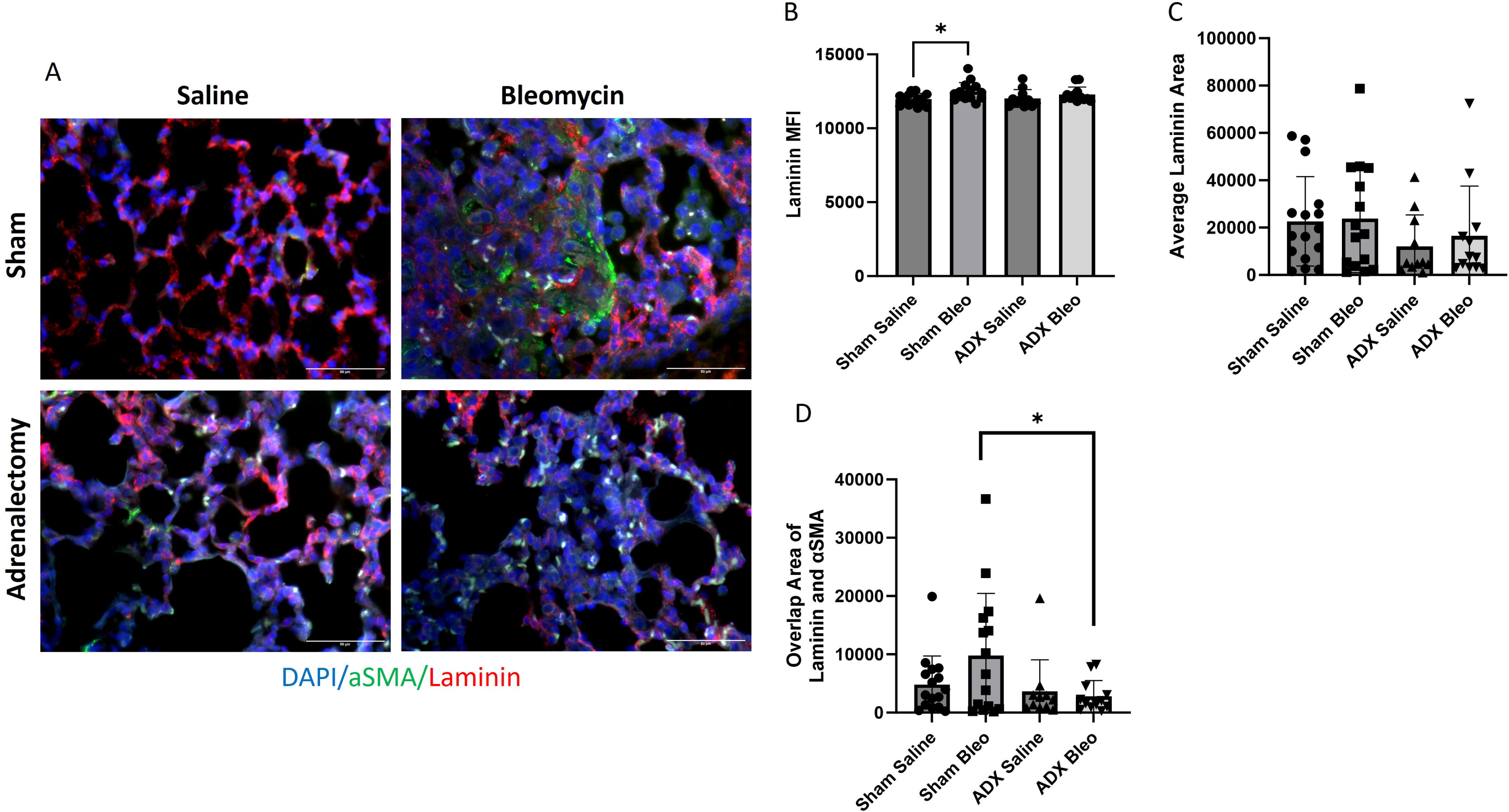
Adrenalectomy impacted the correlation between αSMA and Laminin. Through immunofluorescent imaging, we investigated the alterations in laminin expression and its co-expression with α-smooth muscle actin (αSMA). (A) Immunostaining of Laminin (red) and αSMA (green) in the lung tissue of mice that underwent adrenalectomy or sham surgery in the Bleomycin model of fibrosis. (B) MFI of Laminin was significantly increased by Bleomycin in sham but not adrenalectomy groups. (C) There was no significant change of Laminin positive area among groups. (D) Relative to sham surgery, adrenalectomy significantly reduced the double positive area of αSMA and Laminin. Scale bar = 50 microns. Data are shown as mean +/- SEM. One-way ANOVA was performed to determine significance (* = p<0.05).

**Figure 7.**
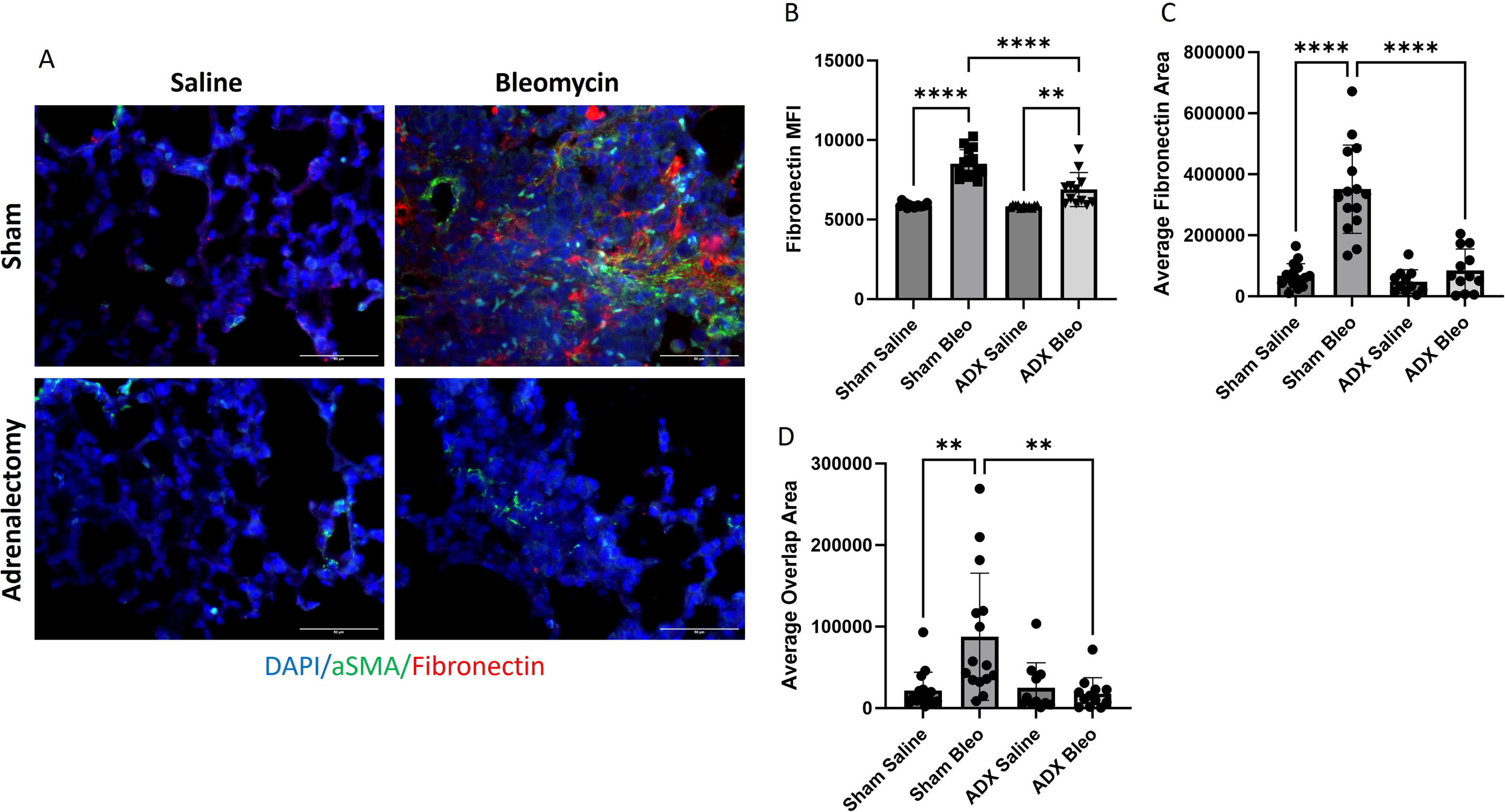
Adrenalectomy reduced fibronectin expression and impacted the correlation between αSMA and fibronectin. Through immunofluorescent imaging, we investigated the alterations in fibronectin expression and its co-expression with αSMA. (A) Representative images of the immunostaining of fibronectin (red) and αSMA (green) are shown. (B) The MFI of fibronectin increased in the sham-bleomycin group and was significantly less after adrenalectomy. (C) The average area of significantly increased following bleomycin stimulation in the mice undergoing sham surgery. However, adrenalectomy effectively reversed this increase. (D) The average area where fibronectin and αSMA overlapped significantly increased following bleomycin stimulation in the mice undergoing sham surgery, while adrenalectomy effectively nullified this increase. For all images αSMA is in the green, FITC channel, Laminin is red, and nuclei are counterstained with DAPI. Scale bar = 50 microns. Data are shown as mean +/- SEM. One-way ANOVA was performed to determine significance (** = p<0.01; *** = p<0.005; **** = p<0.0001).

### Adrenalectomy ablates Adrenaline but not Noradrenaline levels

The adrenal glands play a pivotal role in the production and secretion of AD and NA. In contrast to AD which is primarily synthesized in the adrenal medulla, lung NA is primarily derived from tissue-resident post-ganglionic neurons [27, 28]. We hypothesized that the protective effects of adrenalectomy in lung fibrosis would be induced by marked reductions systemic and pulmonary levels of adrenally derived catecholamines. To validate this hypothesis, we first evaluated AD levels in lung homogenate, BAL, and plasma by ELISA and found that AD was significantly reduced after adrenalectomy (Figure 8 A-C). These data confirm the effectiveness of the adrenalectomy in reducing circulating AD levels. We similarly evaluated NA concentrations in lung homogenate, BAL, and plasma. NA levels in lung homogenate and plasma were unaffected by adrenalectomy, consistent with prior data published by our lab [6], but were reduced in the BAL of saline-control mice subjected to adrenalectomy (Figure 9A-C). The impact of this reduction on lung fibrosis remains to be explored.

**Figure 8.**
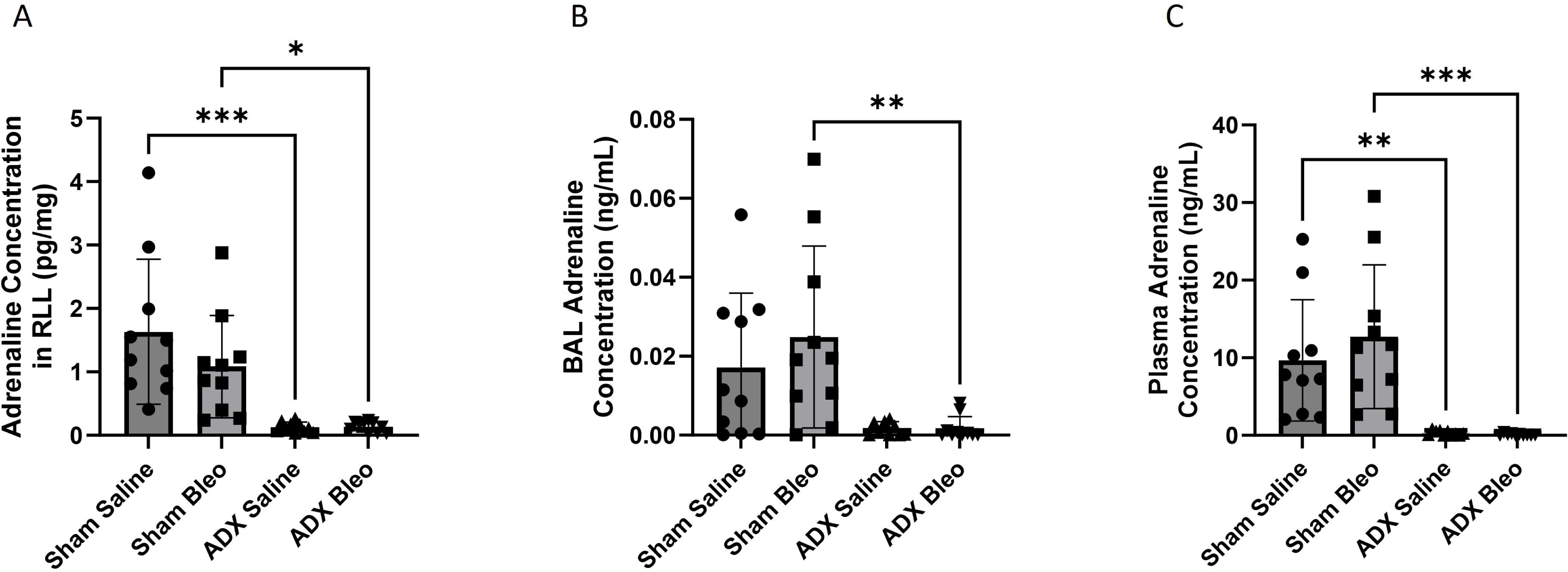
The concentrations of AD in mice underwent sham surgery or adrenalectomy in bleomycin model. The AD concentrations in (A) lung tissues, (B) BAL and (C) plasma are shown. Data are shown as mean +/- SEM. One-way ANOVA was performed to determine significance (* = p<0.05; ** = p<0.01; *** = p<0.005).

**Figure 9.**
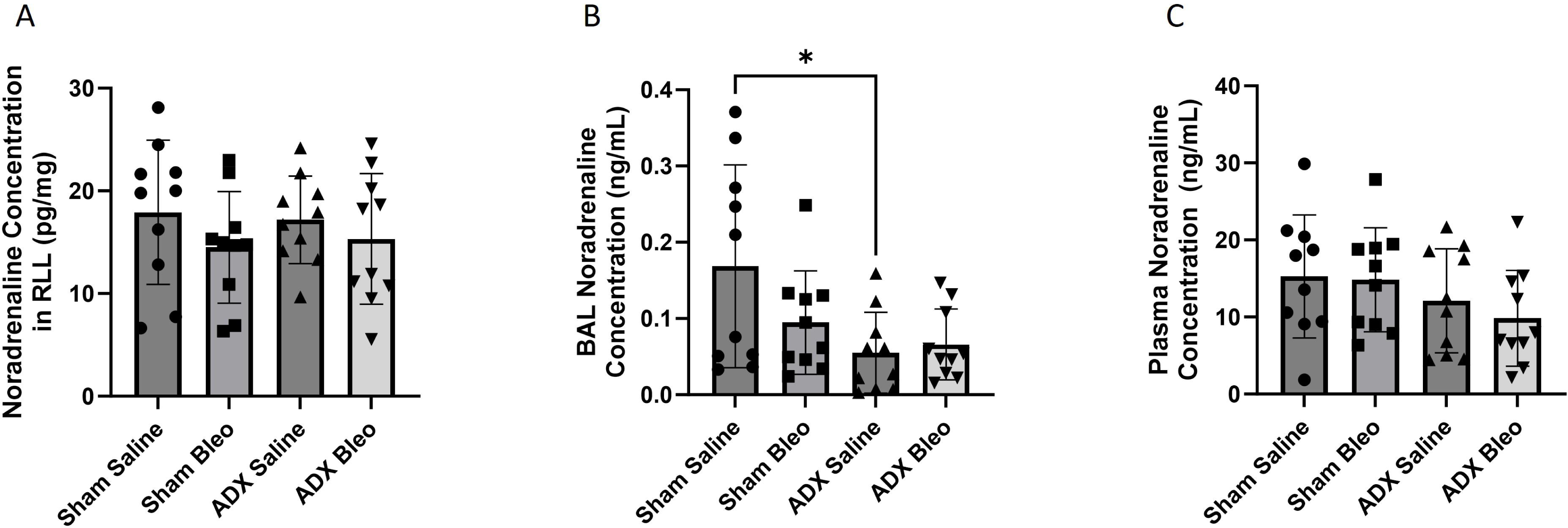
The concentrations of NA in mice underwent sham surgery or adrenalectomy in bleomycin model. The NA concentrations in (A) lung tissues, (B) BAL and (C) plasma are shown. Data are shown as mean +/- SEM. One-way ANOVA was performed to determine significance (* = p<0.05).

### Adrenalectomy eliminates aldosterone production

Aldosterone, secreted by the adrenal glands, is the final common mediator of the RAAS axis [29], and this system has been shown to interact with pro-fibrotic pathways such as TGFβ1 to significantly impact fibrosis in numerous tissues [30, 31]. To gain a more robust understanding of how the processes mediated by the adrenal gland impact pulmonary fibrosis, we measured systemic aldosterone concentration in plasma by ELISA (Figure 10). As expected, we found that the systemic aldosterone concentration was significantly decreased, and even mostly eliminated after adrenalectomy suggesting that RAAS activation in our model was likely decreased or even eliminated and therefore unlikely to contribute to the fibrotic changes induced by bleomycin administration. Nevertheless, further experiments will be needed to better understand the impact of aldosterone on pulmonary fibrosis.

**Figure 10.**
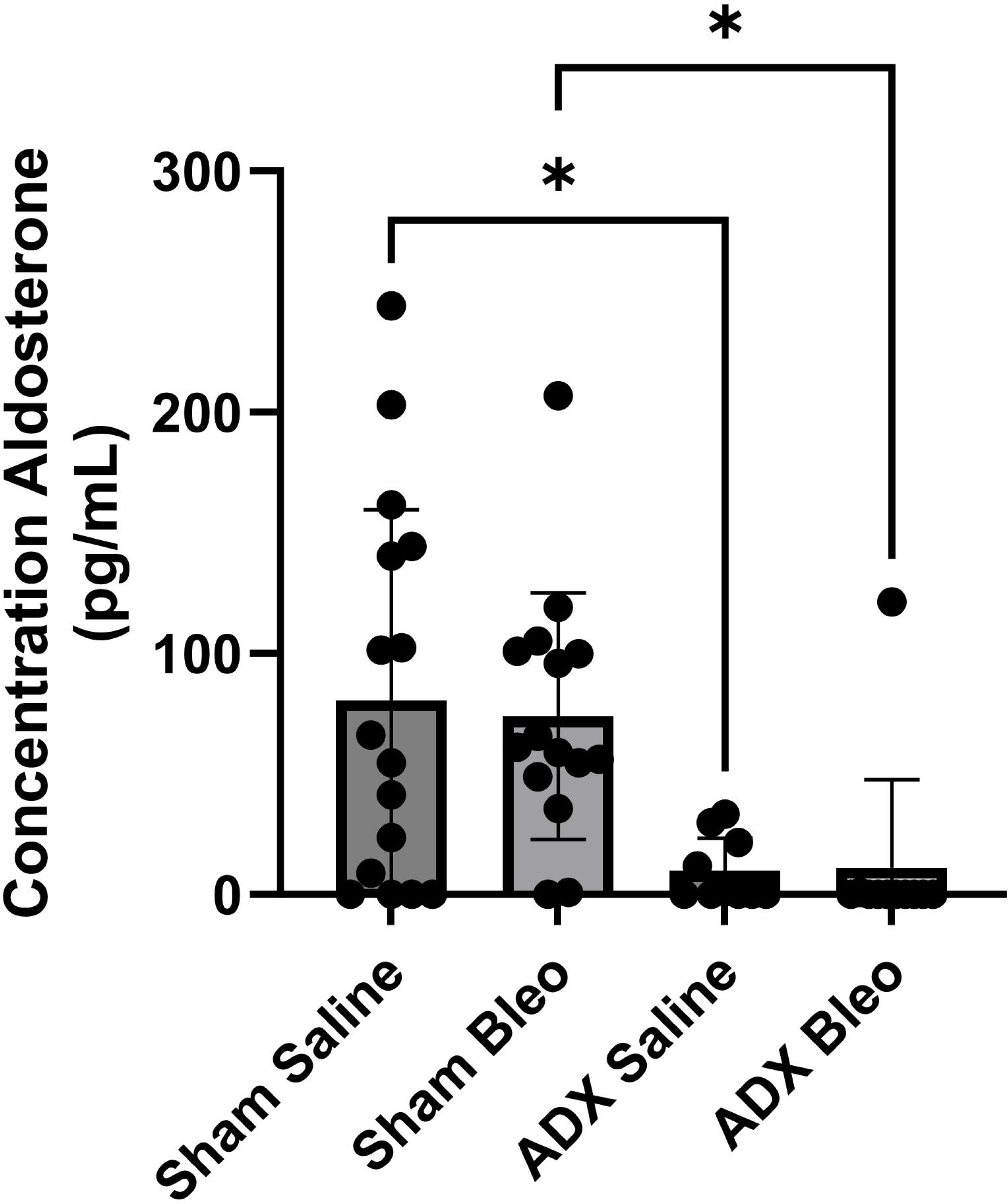
The concentration of aldosterone in mice underwent sham surgery or adrenalectomy in bleomycin model. The concentration of aldosterone in mice underwent sham surgery or adrenalectomy in bleomycin model. Shown are plasma aldosterone concentrations. Data are shown as mean +/- SEM. One-way ANOVA was performed to determine significance (* = p<0.05).

## DISCUSSION

An increasing number of studies have demonstrated catecholamines such as AD and NA associate with fibrosis [6, 8, 32, 33]. However, the contribution of systemically circulating adrenally produced catecholamines to pulmonary fibrosis is less well understood. Here, we investigated the role of surgical adrenalectomy in pulmonary fibrosis using the bleomycin model of experimental lung fibrosis. We found that relative to sham surgery, adrenalectomy significantly decreased airway inflammation by BAL WBC counts. While adrenalectomy increased collagen concentration in uninjured mice, there was no discernible difference in bleomycin induced collagen accumulation. Interestingly, histologic indices of bleomycin induced lung remodeling were reduced by adrenalectomy based on Trichrome-stained images and MAS data. These findings were accompanied by concomitant reductions in αSMA and fibronectin, and amelioration of bleomycin-induced increases of laminin. Whereas adrenalectomy abrogated AD detection in all conditions and compartments, it only reduced BAL NA in uninjured mice. In addition, the systemic aldosterone concentration was significantly decreased.

As expected, airway inflammation, which has been associated with pulmonary fibrosis [34], increased significantly after bleomycin administration and was significantly, though not completely, ameliorated by adrenalectomy. These data suggest that circulating adrenally derived substances impact lung fibrosis, at least in part, by altering inflammatory responses in the lung. Similarly, we noted that adrenalectomy improved but did not completely rescue the lung from histologically assessed fibrosis in the bleomycin model. Surprisingly, adrenalectomy resulted in a significant increase in soluble and insoluble collagen in mice treated saline control but there was no difference observed in mice that received bleomycin (Figure 4). This suggests that the protective effects of adrenalectomy are not related to collagen secretion and deposition and implicates adrenally derived substances as regulators of ECM remodeling. Indeed, while both NA and ALD have be reported to promote fibrosis [6, 8, 29–31], AD has been shown to exhibit antifibrotic effects [32, 33]. In this way, adrenalectomy can result in opposing effects in the lung by improving histologic markers of fibrosis but contributing to increased collagen deposition. While further investigation is needed to better understand these relationships, and though total adrenal ablation may not be an effective treatment option, our data highlight the importance of a lung-adrenal axis in pulmonary fibrosis.

αSMA and ⍺SMA-expressing myofibroblasts are well-known to be important in the development of lung fibrosis [22, 35]. Consistent with previous publications [6, 36], we noted that bleomycin significantly increased the expression of αSMA but that this increase was reversed by adrenalectomy. As αSMA is the principal biomarker of myofibroblasts, our observation suggests that adrenalectomy may result in reduced differentiation of cells into ⍺SMA-expressing myofibroblasts thereby offering some protection against bleomycin-induced fibrosis. Because collagen content did not change between sham and adrenalectomy groups that received bleomycin, we reasoned that the histologic changes we observed must come from other components of the ECM. We thus tested two core ECM proteins, laminin and fibronectin, that are associated with fibrosis [23–26, 37] and found that laminin MFI significantly increased in bleomycin-treated mice that underwent sham surgery but not adrenalectomy. This indicates that laminin expression is increased in mice that received bleomycin, consistent with prior data [23], and suggests that adrenalectomy may be protective against increases in laminin expression. In addition, the co-expression of laminin and αSMA was significantly reduced by adrenalectomy in bleomycin-treated mice versus to sham surgery. Similarly, the expression of fibronectin and its co-expression with αSMA were also increased in bleomycin-treated animals and was significantly abrogated by adrenalectomy. Taken together, our results indicate that adrenally derived substances are important mediators of fibrotic lung remodeling and show that adrenalectomy protects against the development of fibrosis by ameliorating increases in αSMA and critical ECM components. Further investigation into these mechanisms could yield more targeted treatments for pulmonary fibrosis.

Our interest in the functions of the adrenal gland in relation to lung fibrosis began with understanding how AD and NA contribute to the progression of fibrosis. Because of this, it was important to study how the concentrations of these two hormones change between the groups. As expected, we found that AD was almost completely eliminated by adrenalectomy consistent with results seen in our previous experiments [8]. Interestingly, there was almost no significant difference between the NA concentrations of the various groups tested indicating that most of the NA seen is nerve-derived and thus not effected by adrenalectomy. This is consistent with previous findings and implies that targeting locally-derived NA is more likely to benefit fibrotic endpoints [8].

We measured the plasma concentration of aldosterone to take a preliminary look at the activity of the RAAS axis and found that, as expected, there was no systemic aldosterone present after adrenalectomy. In addition to the ECM remodeling effects noted above, elimination of plasma ALD secretion may provide another explanation for the protective effects of adrenalectomy in the bleomycin model. Previous research has demonstrated that RAAS antagonists are protective against bleomycin induced lung injury in patients receiving bleomycin chemotherapy [38] and angiotensin-converting enzyme inhibitors have been shown to be effective in the mitigation of fibrosis caused by the SARS-CoV-2 virus [39]. While our results are promising, further research is needed to understand the interactions between RAAS activity and pulmonary fibrosis.

In conclusion, this study implicates the adrenal gland as an important regulator of pulmonary fibrosis. Mice that underwent adrenalectomy were more resistant to the harmful effects of bleomycin and exhibited changes in ECM deposition and orientation. However, there are significant drawbacks. Adrenalectomy is not a viable treatment modality for patients with pulmonary fibrosis. Additionally, by increasing collagen content, total adrenalectomy clearly has opposing effects on fibrotic remodeling that will require additional study to clarify. Future studies should also evaluate other elements of the ECM such as elastin and the different collagens to gain a better understanding of the specific effects of adrenalectomy. A better understanding of the biology underlying adrenalectomy’s protective effects in pulmonary fibrosis will be fundamental to translating these promising results into effective therapeutic strategies for patients suffering from this devastating disease.

## ACKNOWLEDGEMENTS

This work was supported by The Assistant Secretary of Defense for Health Affairs endorsed by the Department of Defense, in the amount of ($334,999.00), through the Peer Reviewed Medical Research Program under Award Number W81XWH-20-1-0157. Opinions, interpretations, conclusions, and recommendations contained herein are those of the author(s) and are not necessarily endorsed by the Department of Defense. The U.S. Army Medical Research Acquisition Activity, 820 Chandler Street, Fort Detrick MD 21702-5014 is the awarding and administering acquisition office. In conducting research using animals, the investigator(s) adhere(s) to the laws of the United States and regulations of the Department of Agriculture. In the conduct of research involving hazardous organisms or toxins, the investigator(s) adhered to the CDC-NIH Guide for Biosafety in Microbiological and Biomedical Laboratories.

## GRANTS

HS was supported by U.S. Department of Defense (DOD) Award W81XWH-20-1-0157, ATS foundation and Scleroderma Foundation. ARG was supported by 5T32HL007778-27/28 from the NIH. ELH was supported by R01HL152677, and R01HL163984 from the NIH, the Gabriel and Alma Elias Research Fund, and the Greenfield Foundation. The content is solely the responsibility of the authors and does not necessarily represent the official views of the National Institutes of Health or the Department of Defense.

## AUTHOR CONTRIBUTIONS

J.M., C.P., E.L.H. and H.S. conceived and designed research; J. M., C.P., A.G. and H.S. performed experiments; J.M., C.P., A.G. S.S. and H.S. analyzed data; J.M., C.P., A.G., E.L.H. and H.S. interpreted results of experiments; J.M., C.P. and H.S. prepared figures; J.M., C.P., A.G. and H.S. drafted manuscript; J.M., C.P., A.G., S.S., E.L.H. and H.S. edited and revised manuscript; J.M., C.P., A.G., S.S., E.L.H. and H.S. approved final version of manuscript.

## Notes

### Competing Interest Statement

The authors have declared no competing interest.

## REFERENCES

1. Martinez FJ, Collard HR, Pardo A, Raghu G, Richeldi L, Selman M, et al. Idiopathic pulmonary fibrosis. Nat Rev Dis Primers. 2017;3:17074. Epub 2017/10/21. doi: 10.1038/nrdp.2017.74. PubMed PMID: 29052582.

2. Blaauboer ME, Boeijen FR, Emson CL, Turner SM, Zandieh-Doulabi B, Hanemaaijer R, et al. Extracellular matrix proteins: a positive feedback loop in lung fibrosis? Matrix Biology. 2014;34:170–8.

3. Xia Y, Wang H, Shao M, Liu X, Sun F. MAP3K19 Promotes the Progression of Tuberculosis-Induced Pulmonary Fibrosis Through Activation of the TGF-β/Smad2 Signaling Pathway. Molecular Biotechnology. 2023:1–11.

4. Zhao W, Wang L, Wang Y, Yuan H, Zhao M, Lian H, et al. Injured Endothelial Cell: A Risk Factor for Pulmonary Fibrosis. International Journal of Molecular Sciences. 2023;24(10):8749.

5. Phan THG, Paliogiannis P, Nasrallah GK, Giordo R, Eid AH, Fois AG, et al. Emerging cellular and molecular determinants of idiopathic pulmonary fibrosis. 2021;78:2031–57.

6. Gao R, Peng X, Perry C, Sun H, Ntokou A, Ryu C, et al. Macrophage-derived netrin-1 drives adrenergic nerve-associated lung fibrosis. J Clin Invest. 2021;131(1). Epub 2021/01/05. doi: 10.1172/JCI136542. PubMed PMID: 33393489; PubMed Central PMCID: PMCPMC7773383.

7. Berg RA, Moss J, Baum BJ, Crystal RGJTJoCI. Regulation of collagen production by the β-adrenergic system. 1981;67(5):1457–62.

8. Ishikawa G, Peng X, McGovern J, Woo S, Perry C, Liu A, et al. α1 Adrenoreceptor antagonism mitigates extracellular mitochondrial DNA accumulation in lung fibrosis models and in patients with idiopathic pulmonary fibrosis. American Journal of Physiology-Lung Cellular and Molecular Physiology. 2023;324(5):L639–L51. doi: 10.1152/ajplung.00119.2022. PubMed PMID: 36648147.

9. Rassler B, Marx G, Schierle K, Zimmer H-GJCP, Biochemistry. Catecholamines can induce pulmonary remodeling in rats. 2012;30(5):1134–47.

10. Skurikhin EG, Pershina OV, Reztsova AM, Ermakova NN, Khmelevskaya ES, Krupin VA, et al. Modulation of bleomycin-induced lung fibrosis by pegylated hyaluronidase and dopamine receptor antagonist in mice. 2015;10(4):e0125065.

11. Yu AQ, Wang J, Zhou XJ, Chen KY, Cao YD, Wang ZX, et al. Senescent cell-secreted netrin-1 modulates aging-related disorders by recruiting sympathetic fibers. Frontiers in Aging Neuroscience. 2020;12:507140.

12. Gupta D, Kumar A, Mandloi A, Shenoy V. Renin angiotensin aldosterone system in pulmonary fibrosis: Pathogenesis to therapeutic possibilities. Pharmacological Research. 2021;174:105924.

13. Mehdizadeh S, Taherian M, Bayati P, Mousavizadeh K, Pashangzadeh S, Anisian A, et al. Plumbagin attenuates Bleomycin-induced lung fibrosis in mice. 2022;18(1):93.

14. Hubner RH, Gitter W, El Mokhtari NE, Mathiak M, Both M, Bolte H, et al. Standardized quantification of pulmonary fibrosis in histological samples. Biotechniques. 2008;44(4):507–11, 14-7. Epub 2008/05/15. doi: 10.2144/000112729. PubMed PMID: 18476815.

15. Cordelieres FP, Bolte S, editors. JACoP v2. 0: improving the user experience with co-localization studies. ImageJ User & Developer Conference; 2008.

16. Chrousos GPJNEJoM. The hypothalamic–pituitary–adrenal axis and immune-mediated inflammation. 1995;332(20):1351-63.

17. Griton M, Konsman JPJCAR. Neural pathways involved in infection-induced inflammation: recent insights and clinical implications. 2018;28:289–99.

18. Luqman A, Götz FJIJoMS. The ambivalent role of skin microbiota and adrenaline in wound healing and the interplay between them. 2021;22(9):4996.

19. Gilbert KC, Brown NJJCoie, diabetes,, obesity. Aldosterone and inflammation. 2010;17(3):199.

20. Izbicki G, Segel M, Christensen T, Conner M, Breuer RJIjoep. Time course of bleomycin-induced lung fibrosis. 2002;83(3):111–9.

21. Lin Y-H, Wu X-M, Lee H-H, Lee J-K, Liu Y-C, Chang H-W, et al. Adrenalectomy reverses myocardial fibrosis in patients with primary aldosteronism. 2012;30(8):1606–13.

22. Phan SHJPotATS. Genesis of the myofibroblast in lung injury and fibrosis. 2012;9(3):148–52.

23. Lee C-M, Cho SJ, Cho W-K, Park JW, Lee J-H, Choi AM, et al. Laminin α1 is a genetic modifier of TGF-β1–stimulated pulmonary fibrosis. 2018;3(18).

24. Mak KM, Mei RJTAR. Basement membrane type IV collagen and laminin: an overview of their biology and value as fibrosis biomarkers of liver disease. 2017;300(8):1371–90.

25. Philp CJ, Siebeke I, Clements D, Miller S, Habgood A, John AE, et al. Extracellular matrix cross-linking enhances fibroblast growth and protects against matrix proteolysis in lung fibrosis. 2018;58(5):594–603.

26. Muro AF, Moretti FA, Moore BB, Yan M, Atrasz RG, Wilke CA, et al. An essential role for fibronectin extra type III domain A in pulmonary fibrosis. 2008;177(6):638–45.

27. Webster R. Neurotransmitters, drugs and brain function: John Wiley & Sons; 2001.

28. Estelius J. Neuroimmune mechanisms in chronic inflammation: translational studies of the inflammatory reflex: Karolinska Institutet (Sweden); 2018.

29. Azizan EA, Drake WM, Brown MJ. Primary aldosteronism: molecular medicine meets public health. Nature Reviews Nephrology. 2023;19(12):788–806.

30. Brown NJJNRN. Contribution of aldosterone to cardiovascular and renal inflammation and fibrosis. 2013;9(8):459–69.

31. Azibani F, Fazal L, Chatziantoniou C, Samuel J-L, Delcayre CJChr. Aldosterone mediates cardiac fibrosis in the setting of hypertension. 2013;15:395–400.

32. Thong KX, Andriesei P, Luo J, Qin M, Ng J, Tagalakis AD, et al. Adrenaline blocks key cell cycle genes and exhibits antifibrotic and vasoconstrictor effects in glaucoma surgery. 2023;233:109561.

33. Song L, Tian Y, Xu Z-J, Zhang C-P. Adrenaline inhibited cell proliferation and regulated expression of TGF-beta1 and bFGF in cultured human hypertrophic scar fibroblasts via alpha-receptor. 2008.

34. Van Hoecke L, Job ER, Saelens X, Roose K. Bronchoalveolar lavage of murine lungs to analyze inflammatory cell infiltration. JoVE (Journal of Visualized Experiments). 2017;(123):e55398.

35. Biasin V, Crnkovic S, Sahu-Osen A, Birnhuber A, El Agha E, Sinn K, et al. PDGFRα and αSMA mark two distinct mesenchymal cell populations involved in parenchymal and vascular remodeling in pulmonary fibrosis. 2020;318(4):L684–L97.

36. Higashiyama H, Yoshimoto D, Kaise T, Matsubara S, Fujiwara M, Kikkawa H, et al. Inhibition of activin receptor-like kinase 5 attenuates bleomycin-induced pulmonary fibrosis. 2007;83(1):39–46.

37. Savin IA, Zenkova MA, Sen’kova AV. Pulmonary fibrosis as a result of acute lung inflammation: Molecular mechanisms, relevant in vivo models, prognostic and therapeutic approaches. International Journal of Molecular Sciences. 2022;23(23):14959.

38. Hara R, Onizuka M, Shiraiwa S, Harada K, Aoyama Y, Ogiya D, et al. The role of hypertension and renin-angiotensin-aldosterone system inhibitors in bleomycin-induced lung injury. Clinical Lymphoma Myeloma and Leukemia. 2021;21(4):e321–e7.

39. Elshafei A, Khidr EG, El-Husseiny AA, Gomaa MH. RAAS, ACE2 and COVID-19; a mechanistic review. Saudi journal of biological sciences. 2021;28(11):6465–70.

